# Repertoire, function, and structure of serological antibodies induced by the R21/Matrix-M malaria vaccine

**DOI:** 10.1101/2024.10.07.617084

**Authors:** Jonathan R. McDaniel, William N. Voss, Georgina Bowyer, Scott A. Rush, Alexandra J. Spencer, Duncan Bellamy, Marta Ulaszewska, Jule Goike, Scott Gregory, C. Richter King, Jason S. McLellan, Adrian V.S. Hill, George Georgiou, Katie J. Ewer, Gregory C. Ippolito

## Abstract

The World Health Organization recently recommended the programmatic use of R21/Matrix-M vaccine for *Plasmodium falciparum* malaria prevention in children living in malaria-endemic areas. To determine its effects on humoral immunity, we conducted a proteomic analysis of polyclonal IgG antibodies directed against the NANP tetrapeptide of the circumsporozoite protein (CSP) which comprises the vaccine’s core immunogen. In ten malaria-naïve adult volunteers, R21/Matrix-M induced polarized IgG anti-NANP repertoires, heavily skewed for *IGHV3-30/3-33* genes bearing minimal somatic mutation, which remained static in composition following a controlled human malaria infection challenge. Notably, these vaccine-generated antibodies cross-reacted with another protective CSP epitope, the N-terminal junction region, despite its absence from the R21 construct. NANP-specific *IGHV3-30/3-33* monoclonal antibodies mined from polyclonal IgG repertoires blocked sporozoite invasion *in vitro* and prevented parasitemia *in vivo*. Overall, R21/Matrix-M elicits polarized, minimally mutated, polyclonal IgG responses that can target multiple protective CSP epitopes, offering molecular insight into the serological basis for its demonstrated efficacy against *P. falciparum* malaria.

## INTRODUCTION

*Plasmodium falciparum* (Pf) malaria is one of the world’s most ancient diseases and remains the deadliest parasitic disease worldwide. The World Health Organization (WHO) estimated there were approximately 608,000 malaria-related deaths worldwide in 2022 (Venkatesan, 2024). Although there are vaccine candidates under development that target each stage of the parasite’s life cycle, vaccines targeting the pre-erythrocytic malarial sporozoite phase are the most advanced. This malaria vaccine approach remains an area of intense research due to its potential to entirely circumvent liver and blood-stage infection, and thereby prevent disease and onward transmission.

The circumsporozoite protein (CSP), the primary outer coating essential for sporozoite development and hepatocyte invasion, has long been an attractive malaria vaccine target as it is highly immunogenic and capable of inducing a dominant neutralizing titer after infection or vaccination (Hill, 2006; Kappe et al., 2004; Nussenzweig and Nussenzweig, 1989). Earlier *in vitro* studies have demonstrated antibodies targeting CSP can prevent sporozoites from infecting liver cells (Hollingdale et al., 1984). In murine models, both passive transfer of polyclonal and monoclonal antibodies against the immunodominant B-cell major repeat epitope of CSP and active immunization with constructs containing this epitope confer protection against sporozoite challenge (Foquet et al., 2014; Oyen et al., 2017; Wang et al., 1995; Zavala and Chai, 1990).

*P. falciparum* CSP (PfCSP) consists of three antigenic domains: an N-terminal junctional domain with a highly conserved pentapeptide sequence, an immunodominant central repeat region consisting of ∼45 tetrapeptides (NANP “major repeat” and NPDP and NVDP “minor repeats”) and concludes with a C-terminal domain. The Asn-Ala-Asn-Pro (NANP) tetrapeptide of the central repeat region constitutes the immunodominant B-cell epitope of PfCSP (Dame et al., 1984; Zavala et al., 1983), and immunoglobulin G1 (IgG1) and IgG3 constitute the majority of the anti-PfCSP serological response (Ubillos et al., 2018). After more than six decades of diligent effort to create an effective pre-erythrocytic malaria vaccine based on PfCSP, in the last three years, two have been recommended by WHO for use in children living in malaria-endemic areas: RTS,S/AS01 (brand name Mosquirix, by GlaxoSmithKline) and R21/Matrix-M (developed by the University of Oxford’s Jenner Institute and manufactured by the Serum Institute of India).

RTS,S/AS01, recommended by WHO in 2021, is a chimera between the hepatitis B surface antigen (HBsAg) and a truncated form of PfCSP that includes a portion of the unstructured and immunodominant NANP repeating peptide and the carboxy-terminus domain (Miura et al., 2024; White et al., 2015; Zavala, 2022). Although several recent clinical trials have demonstrated RTS,S is safe and offers a high degree of protection immediately following vaccination, both the anti-CSP antibody titer and vaccine efficacy against infection wane over time (White et al., 2015; Zavala, 2022).

R21/Matrix-M was added to the WHO recommendation in October 2023 due to its demonstrated comparable efficacy to RTS,S/AS01 (Datoo et al., 2024; Moorthy et al., 2024; World Health Organization, 2024). The availability of R21/Matrix-M complements the ongoing distribution of RTS,S (Nnaji et al., 2024; Schmit et al., 2024) and boosts overall access to malaria vaccines. R21/Matrix-M is a virus-like particle (VLP) vaccine which targets the same CSP antigen as RTS,S and consists of *P. falciparum* strain NF54, containing 19 copies of the NANP central repeat, and is formulated with Matrix-M adjuvant. Clinical trials of R21/Matrix-M have demonstrated high vaccine efficacy over one year with a three-dose regimen in 4,644 children aged 5 to 36 months. Results indicate efficacy rates of 75% (95% CI 71–79) at sites with moderate to high seasonal malaria transmission and 68% (95% CI 61–74) at sites with low to moderate perennial malaria transmission (Datoo et al., 2024), which is comparable to RTS,S/AS01 in similar settings (Chandramohan et al., 2021; RTS, 2015). At this stage, trials of R21/Matrix-M have not been conducted in areas of high perennial transmission, and published trials do not yet allow for direct comparisons between efficacy of the two vaccines. Mathematical modeling of three years of follow-up data stemming from an earlier Phase 2b trial found that anti-CSP antibody titers satisfy the criteria for an immunological correlate of protection for R21/Matrix-M vaccine efficacy against clinical malaria (Schmit et al., 2024).

Whereas prior studies have examined the molecular features of anti-CSP antibody responses when elicited by natural infection, attenuated Pf whole-sporozoite (PfSPZ) vaccination, or RTS,S/AS01 vaccination, the molecular features of anti-CSP antibody responses following R21/Matrix-M vaccination have yet to be investigated. Until now, human CSP-reactive monoclonal antibodies (mAbs) have been isolated exclusively by single-cell cloning using peripheral B cells (Imkeller et al., 2018; Kisalu et al., 2018; Murugan et al., 2018; Oyen et al., 2017; Tan et al., 2018; Triller et al., 2017; Williams et al., 2024). In these studies, B cells express germline antibodies or antibodies with low levels of mutation that preferentially express the highly homologous, allelically diverse *IGHV3-30* or *IGHV3-33* gene segments, which can mediate NANP recognition via a conserved tryptophan residue at position 52 (Trp52) of the germline-encoded complementarity determining region 2 of the heavy-chain variable region (CDR-H2) (Pholcharee et al., 2020). In contrast, the CSP epitopes targeted by the actual circulating plasma immunoglobulins (IgG) in vaccinated study subjects—especially those plasma antibodies that are most abundant and thereby play a dominant role in the polyclonal response—remain unknown.

Technological advances now provide a means to directly probe endogenous plasma IgG antibodies (de Graaf et al., 2022; Lavinder et al., 2015; Schulte et al., 2022; Townsend et al., 2024). Here, capitalizing on such advances, we set out to address at the molecular-level the PfCSP humoral immune response induced by R21/Matrix-M vaccination in malaria-naïve volunteers, analyzing the plasma IgG antibody response to a NANP_6_ peptide (NANP hexamer) post-vaccination as well as before and after a controlled human malaria infection (CHMI) challenge. In this study, we show that the R21/Matrix-M malaria vaccine elicits a highly focused IgG humoral response in malaria-naïve individuals, predominantly involving minimally mutated antibodies utilizing *IGHV3-30/3-33* genes, consistent with previous observations of B-cell responses. Our findings extend this understanding by demonstrating that the IgG response remains both stable in quantity and unaltered in its antibody composition, even following controlled human malaria infection. Additionally, a key discovery is the cross-reactivity of these vaccine-induced IgG antibodies with the N-terminal junction of the circumsporozoite protein, an epitope not included in the R21 vaccine, indicating epitope spreading within the serological repertoire. Altogether, our data offer a molecular-level dissection of R21/Matrix-M IgG humoral immunity and provide a foundation to explain, in part, the demonstrated vaccine efficacy of R21/Matrix-M.

## RESULTS

### Antibody lineage diversity in R21/Matrix-M vaccinated individuals

A synergistic combination of immunoglobulin mass spectrometry (Ig-Seq) and B-cell receptor deep sequencing (BCR-Seq) were used to analyze the molecular composition and relative abundance of circulating anti-NANP IgG antibodies. (See Methods.) For Ig-Seq proteomics, total IgG purified from volunteer plasma was subjected to affinity chromatography using recombinant NANP polypeptide (Asn-Ala-Asn-Pro hexamer [NANP_6_]) representative of the CSP central repeat region. The eluted fraction (“NANP_6_-reactive” IgG) and flow-through fraction (“NANP_6_-nonreactive” IgG) were analyzed by high-resolution liquid chromatography and tandem mass spectrometry (LC–MS/MS). For BCR-Seq, next-generation sequencing (NGS) of peripheral blood B-cell receptor (BCR) heavy-chain (VH), light-chain (VL), and single B-cell paired VH:VL variable region repertoires was performed. High-confidence IgG peptide spectral matches from the LC-MS/MS could then be identified using these volunteer-specific BCR databases— focusing on VH-derived hypervariable CDR-H3 peptides in particular—to determine the presence and relative abundance of IgG antibodies at both the lineage (family) and clonal (individual) levels. The integration of IgG peptide sequencing (Ig-Seq) with matching VH/VH:VL DNA sequencing (BCR-Seq) (Lavinder et al., 2015; Lavinder et al., 2014; Voss et al., 2021) allowed us to precisely identify and quantify circulating NANP_6_-reactive IgG repertoires across a volunteer cohort at the molecular level. This approach also facilitated the cloning and validation of a small panel of recombinant IgG anti-NANP plasma mAbs for subsequent validation and functional assessment.

To evaluate the quality and evolution of the IgG humoral immune response to vaccination, we analyzed peripheral blood mononuclear cells (PBMC) and plasma samples collected from ten malaria-naïve volunteers in two Phase I/IIa clinical trials, VAC053 (n=4) and VAC065 (n=6) (**Fig. 1 A**). In both trials, three intramuscular vaccinations were administered in four-week intervals on days 0, 28, and 56. Antibody-secreting plasmablasts were collected from all volunteers one week after the third and final vaccination (day 63). For VAC053, post-vaccination plasma was collected for analysis at day 63. For the VAC065 trial, volunteers were challenged by CHMI with live sporozoites four weeks after the day 56 final vaccination (Day 84), with post-vaccination pre-challenge plasma collected on day 84 (referred to as “C_-1_”) and then monitored for protection from infection until a final blood draw five weeks later at day 112 (referred to as “C_+35_”). Thus, plasma samples were analyzed post-vaccination (VAC053 and VAC065) and post-challenge (VAC065).

**Figure 1:**
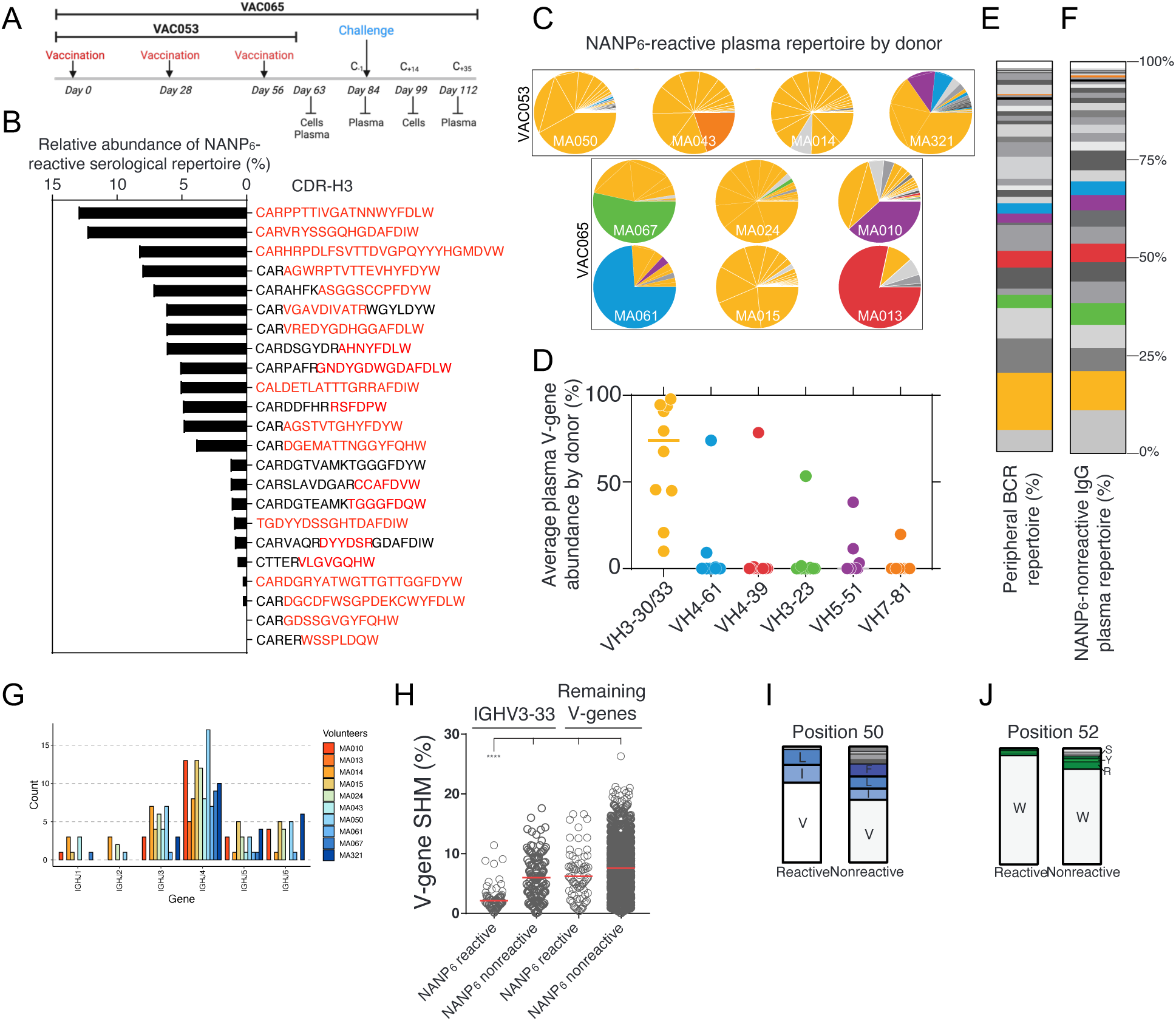
Minimally mutated *IGHV3-30/3-33* genes dominate the IgG anti-NANP_6_ plasma repertoire following three doses of R21/Matrix-M vaccine. (A) Schematic of the VAC053 and VAC065 clinical trials. Plasma and cells (PBMCs) were collected at the times indicated. Volunteers (n=10) were vaccinated at three time points (days 0, 28, 56). Ig-Seq analyses performed at day 63 (VAC053) or day 84 (VAC065). **(B)** Relative abundance of NANP_6_-reactive IgG lineages in a representative volunteer. Each bar represents a single IgG lineage identified by its respective CDR-H3. Red indicates the unique peptides for each lineage that were detected by LC-MS/MS. **(C)** Relative plasma antibody IGHV-gene segment usage for each volunteer. IGHV-gene identity by color is shown in panel D. **(D)** Average IGHV-gene abundance in NANP_6_-reactive IgG. Bar represents mean. **(E)** IGHV-gene frequency, by lineage, in total peripheral B cells averaged across all volunteers. **(F)** IGHV-gene frequency, by lineage, in NANP_6_-nonreactive IgG averaged across all volunteers. **(G)** IGHJ gene usage in same volunteer as panel B. **(H)** IGHV-gene somatic hypermutation levels for NANP_6_-reactive and nonreactive plasma IgG antibodies and B-cell receptor sequences grouped by lineage. ****P<0.0001 (two-tailed Kruskal-Wallis test). Amino acid usage in CDR-H2 **(I)** Position 50 and **(J)** Position 52 of *IGHV3-33* plasma antibodies separated by NANP_6_ reactivity. I = isoleucine; L = leucine; V = valine; W = tryptophan.

First, we set out to address fundamental questions about IgG repertoires generated by R21/Matrix-M vaccination: Are they simple or complex in terms of the number of antibody lineages (i.e., families of closely related antibody VH sequences)? Is the distribution of lineages uniform or polarized? In our study the average IgG anti-NANP_6_ titer among the ten volunteers was 38.5 ± 20.3 µg/mL, as determined by ELISA (**Table 1**). This result indicated a substantial level of circulating IgG, which was sufficient for further analysis using NANP_6_ affinity chromatography and Ig-Seq proteomic deconvolution to identify and quantify IgG lineages. Because this was a “virgin” immune response in malaria-naïve volunteers, these IgG lineages likely derived from the activation of a single naïve B cell which then clonally diversified through somatic hypermutation (SHM).

**Table 1.**
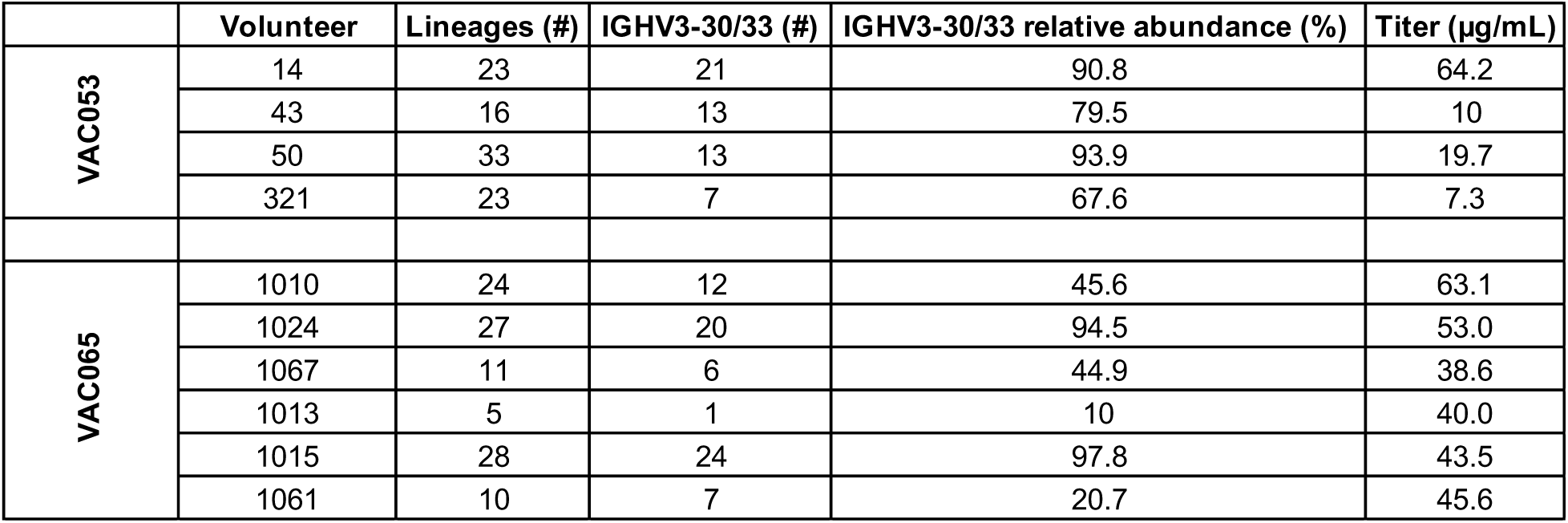
Summary of VAC053 and VAC065 clinical trials and the IgG humoral immune response to R21/Matirx-M vaccination. Titers against NANP_6_ peptide measured by indirect ELISA at day 63 (VAC053) and day 84 (VAC065).

In one representative volunteer, the IgG anti-NANP_6_ repertoire comprised 21 distinct antibody lineages, each exceeding a relative abundance threshold >0.5% (**Fig. 1 B**). Similarly, the size of the IgG anti-NANP_6_ repertoire was consistent across VAC053 and VAC065 volunteers, with an average of 20 ± 9 antibody lineages (range: 5–33) contributing to the NANP_6_ reactivity (**Fig. S1 A, Table 1**, and **Table S1**), similar to other reports of antigen-specific serological repertoires analyzed through proteomic mass spectrometry (Bondt et al., 2021; Townsend et al., 2024). The serological IgG anti-NANP repertoires also displayed a significant degree of polarization; typically, the 2-5 most prevalent lineages, out of the approximately 20 per volunteer, accounted for over 50% of the NANP_6_ antigen-reactive titer (**Fig. 1 C** and **Table S1**).

Despite IgG repertoires being oligoclonal and polarized, the CDR-H3 region of antibody combining sites—a somatically generated, key determinant of binding specificity for most antibodies (Ippolito et al., 2006; Xu and Davis, 2000)—was diverse in both length (range: 6–26 aa) (**Fig. S1 B**) and sequence composition (**Table S1**). Among 197 total IgG lineages derived from the ten volunteers, there was no CDR-H3 repertoire overlap, nor any apparent repertoire convergence among volunteers according to non-linear dimensionality reduction (tSNE) analysis, and therefore no evidence for “public” multi-donor anti-NANP_6_ antibodies (**Fig. S1 C and D**).

### Dominant *IGHV3-30/3-33* gene usage in R21/Matrix-M vaccinated individuals

Contrary to these features of diversity, a common characteristic across all the NANP_6_-reactive IgG repertoires was an obvious enrichment for the variable-region gene segments *IGHV3-30*, *IGHV3-33*, or their related alleles (**Fig. 1 D** and **Fig. S1 E**). This VH family, including alleles with serine, arginine, or tryptophan at position 52 of the CDR-H2 (collectively referred to as “*IGHV3-30/33*”), encompasses *IGHV3-30*, *IGHV3-30-3*, *IGHV3-30-5*, and *IGHV3-33* alleles that share over 96% identity (Tan et al., 2018). On average, *IGHV3-30/33* circulating IgG antibodies constituted ∼70% of the NANP_6_-reactive repertoire post-vaccination (**Fig. 1 D**). Similarly, most PfCSP repeat monoclonal antibodies isolated by single-cell cloning of peripheral human B cells are predominantly encoded by *IGHV3-30/33* (Julien and Wardemann, 2019). While other IGHV genes played dominant roles in certain individuals in both the VAC053 and the VAC065 cohort (**Fig. 1 C and D**), their infrequent usage suggests volunteer-specific, non-public motifs. In 4-of-6 VAC065 volunteers non-*IGHV3-30/33* genes increased in relative plasma abundance by day 84 post-vaccination (MA010, MA013, MA061, and MA067); however, this was not the case for 2-of-6 VAC065 volunteers (MA015 and MA024) wherein *IGHV3-30/33* dominance continued to prevail. We then compared the total peripheral B-cell IgG repertoire from next-generation sequencing (**Fig. 1 E**) with the NANP_6_-nonreactive plasma IgG repertoire (**Fig. 1 F**). Averaged over the 10 volunteers, the similarity in *IGHV3-30/33* frequency between the BCR sequencing repertoire (14.7%) and the nonreactive plasma repertoire (10.1%) indicates that the observed ∼70% IGVH3-30/33 bias in the NANP_6_-binding plasma IgG repertoire reported in **Fig. 1 D** must be due to antigen-specific enrichment. In contrast to the IGHV bias, IGHJ gene usage in plasma or the BCR repertoire was highly diversified and non-polarized (**Fig. 1 G**).

Another common characteristic of NANP_6_-reactive plasma antibodies was a significant reduction in somatic hypermutation (SHM) among those using the *IGHV3-33* gene segment. Antibody lineages not using *IGHV3-33* showed SHM rates approximately 3-fold higher than those using *IGHV3-33* (6.2% ± 4.8 vs 2.1% ± 1.9; **Fig. 1 H**). In support of the hypothesis that NANP recognition is fundamentally enhanced by amino acids encoded in the germline of the *IGHV3-33* gene segment (CDR-H2 Val50 and Trp52, in particular) (Wahl and Wardemann, 2022), it was found that NANP_6_-nonreactive *IGHV3-33* antibodies were nearly three times more mutated compared to NANP_6_-reactive antibodies (6.0% ± 3.8 vs 2.1% ± 1.9, respectively). Of note, nonreactive antibodies had a higher number of replacement mutations at positions Val50 and Trp52 (**Fig. 1, I and J**). Altogether, our results illustrate that the antibody features of the serological repertoire are consistent with findings from the analysis of IgG receptors isolated by B-cell cloning showing that *IGHV3-30/33* genes (*IGHV3-33* primarily) confer binding to the CSP NANP major repeat without reliance on any exact CDR-H3 structure, IGHJ sequence, or acquired somatic mutations during B-cell evolution.

### Antibody lineage diversity in R21/Matrix-M vaccinated individuals remains unchanged following sporozoite challenge

Mosquito bites delivering sporozoite-stage malaria parasites have traditionally served as a model for testing pre-erythrocytic stage vaccines. Here, to gain a deeper understanding of the stability and quality of IgG anti-NANP antibodies induced by R21/Matrix-M vaccination, *Anopheles stephensi* mosquitoes and the chloroquine-sensitive NF54 strain (3D7 clone) of *P. falciparum* were used for CHMI. To evaluate the immune response to vaccination and sporozoite challenge, we analyzed the IgG repertoire post-vaccination in n=5 volunteers at day 84 (“C_-1_”) and then monitored for protection from infection until a final blood draw five weeks later at day 112 (“C_+35_”). **Fig. 2A** and **2B** show plasma antibody repertoires induced by R21/Matrix-M vaccination remained stable despite challenge, with a protected (**Fig. 2A**) and a non-protected (**Fig. 2B**) volunteer displaying consistent IgG lineage composition and relative abundance profiles for their NANP_6_-reactive repertoires. The high degree of correlation between pre- and post-challenge repertoires is shown in scatter plots representing the relative abundance of each identified lineage at the time of plasma collection. The relative abundance of detectable NANP_6_-reactive IgG antibodies in protected volunteer MA-1010 (**Fig. 2A**), for instance, was strikingly correlated lineage-for-lineage and nearly indistinguishable pre- and post-challenge (Pearson correlation, r^2^ = 0.98). In non-protected subject MA-1015, the pre- and post-challenge IgG repertoires were similarly highly correlated at r^2^ = 0.88 (**Fig. 2B**). A strong degree of correlation (r^2^ > 0.6) held for all five volunteers (**Fig. S2**). Thirty-five days, or approximately two IgG1 half-lives, separated the pre- and post-challenge time points, suggesting that the volunteers’ static IgG repertoires are maintained by established, dynamic pools of antibody-secreting cells.

**Figure 2:**
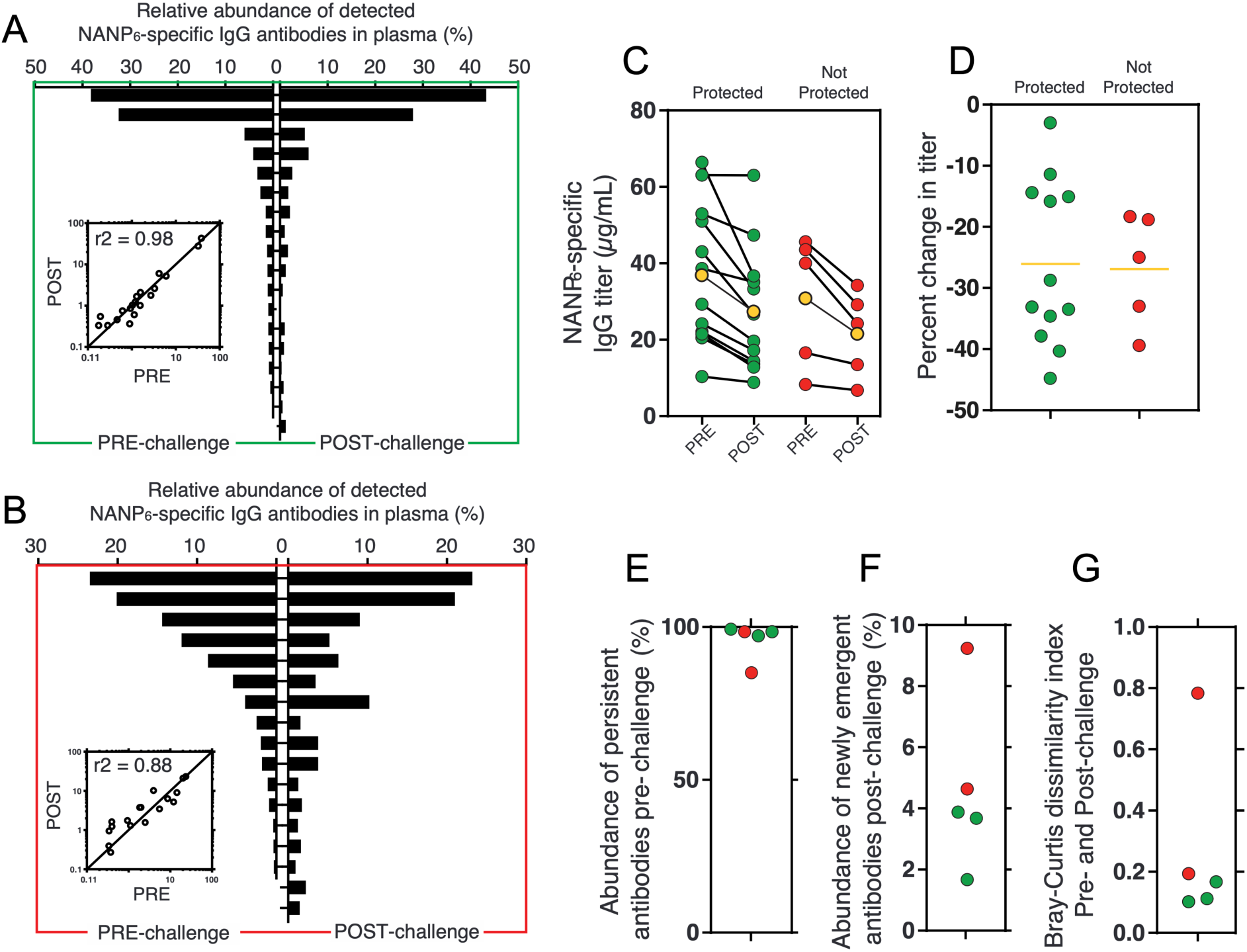
Following vaccination with R21/Matrix M, NANP_6_ reactive plasma antibody repertoires remain highly correlated pre- and post-challenge with live sporozoites. A subset of thrice-vaccinated VAC065 volunteers (n=5) was analyzed at day 84 (pre-challenge; C_-1_) and five weeks later at day 112 (post-challenge; C_+35_). **(A–B)** Relative abundance of NANP_6_ reactive IgG lineages pre- and post-challenge for one representative protected individual (A) and one non-protected individual (B). Each bar represents one lineage. Inset indicates relative abundance for each lineage pre- and post-challenge (Pearson r^2^=0.98 (A) and r^2^=0.88 (B)). **(C)** Estimated plasma titer and **(D)** change in titer of NANP_6_ reactive IgG for volunteers pre- and post-challenge (n=17). Yellow markers indicate the group average. **(E)** Abundance of persistent antibodies pre-challenge (n=5). **(F)** Abundance of emergent antibodies post-challenge (n=5). **(G)** Bray-Curtis dissimilarity index between pre- and post-challenge repertoires (n=5). Lower values indicate higher degrees of similarity. Green indicates protection from infection; red indicates no protection.

Given the static composition of IgG lineages pre- and post-challenge with sporozoites, we wondered whether this molecular feature might also be reflected at the bulk serological level. Using a larger VAC065 cohort of 17 volunteers, we observed NANP_6_ titers in both protected and non-protected groups showed no boosting effect after sporozoite challenge (**Fig. 2C** and **2D**). In fact, collectively, titers declined by an average of 27% between days C_-1_ and C_+35_ over the five-week period between challenge and plasma collection. No significant differences were observed pre-challenge or post-challenge between the absolute (µg/mL) or the relative (%) changes in titer when comparing protected and nonprotected individuals. Upon reexamination of the subset of five volunteers analyzed by Ig-Seq, we note that persistent antibodies, defined as those present pre- and post-challenge, accounted for over 95% of the total NANP_6_-reactive repertoire (**Fig. 2E**), with less than 5% of plasma antibodies post-challenge being detectable only at the second time point (**Fig. 2F**), which could explain in part the static or else decreased titers measured by bulk serology.

### Cross-reactive antibodies induced by R21/Matrix-M target multiple CSP epitopes

PfCSP features a “junction region” (JR) that connects the N-terminal domain to the central NANP repeat region. In the Pf reference isolate 3D7 (PfCSP_3D7), the JR starts with the sequence NPDP, followed by three interspersed NANP and NVDP repeats, with NPDP being unique to this region. Studies have shown that monoclonal antibodies targeting the JR epitope in both mice and humans can be protective (Kayentao et al., 2022; Kayentao et al., 2024; Wang et al., 2020).

B-cell repertoire analyses have identified antibodies specific to the NANP sequence and those cross-reactive with the JR are encoded by *IGHV3-30/33* genes, particularly *IGHV3-33* paired with *IGKV1-5* (reviewed in Julien and Wardemann, 2019). In contrast, antibodies specific to the NVDP sequence and those cross-reactive with both NVDP and NPDP utilize various IGHV gene segments (Kisalu et al., 2018; Murugan et al., 2020). Notably, all JR-binding monoclonal antibodies exhibit some level of cross-reactivity with NANP (Murugan et al., 2020). Despite the R21 vaccine containing only a truncated form of PfCSP with 19 NANP repeats and the C-terminus—lacking the N-terminus, as well as the NPDP and NVDP repeats of the JR (Collins et al., 2017; Stoute et al., 1997)—we found R21/Matrix-M could induce JR-reactive plasma antibodies in all five VAC065 volunteers tested at day 84 post-vaccination (**Fig. 3**). The depletion of pre-challenge (C_-1_) and post-challenge (C_+35_) polyclonal IgG using NANP_6_ affinity chromatography led to a significant reduction in JR-binding signals, as measured by indirect ELISA with a peptide encompassing the JR epitope (junctional region peptide [JR]: KQPADGNPDPNANPNVDPN).

**Figure 3:**
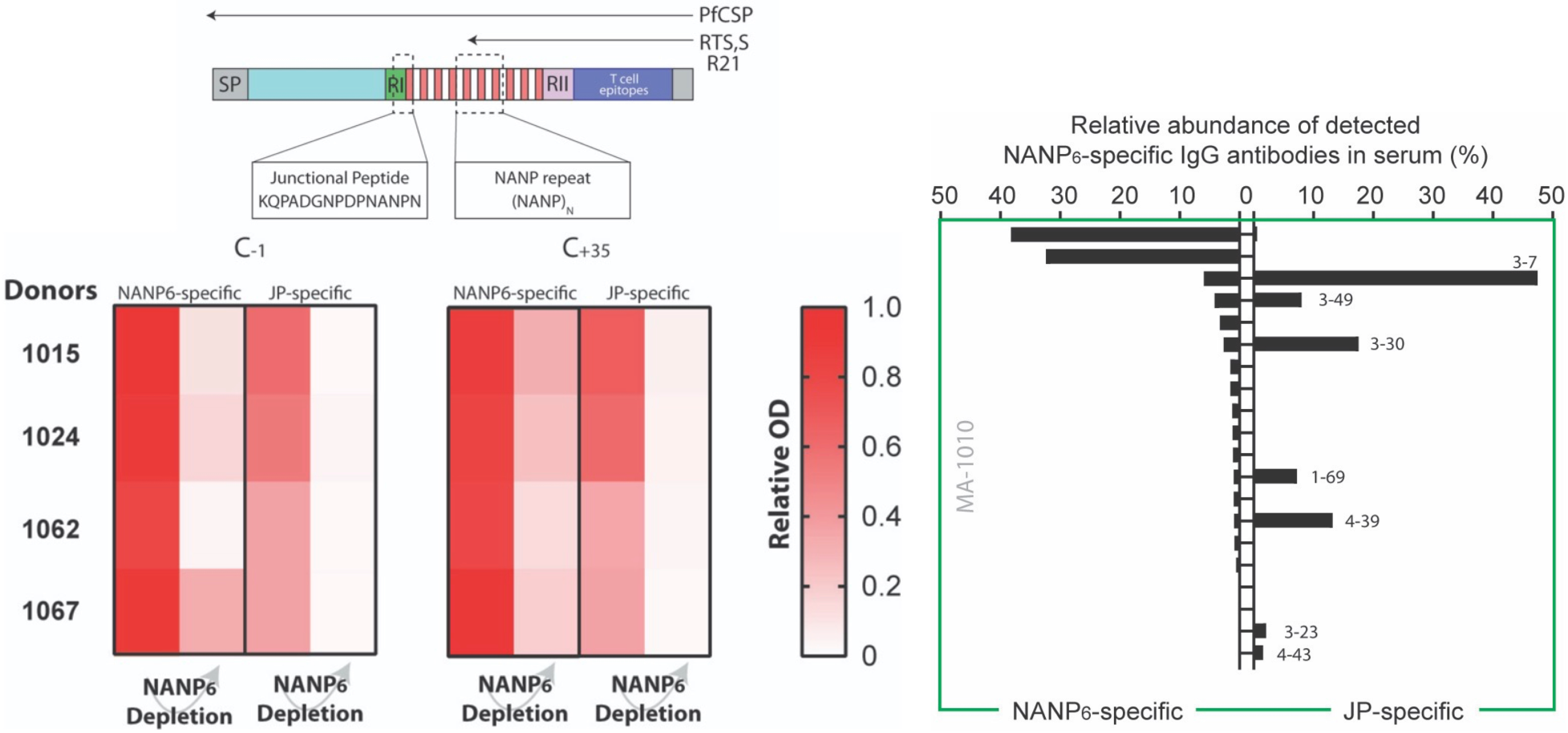
R21/Matrix-M elicits polyclonal IgG plasma antibodies cross-reactive with the N-terminal junction region (JR) of CSP despite its absence in the vaccine construct. NANP_6_ affinity-chromatography depletes the plasma of JR-specific antibodies. The schematic illustrates the truncated CSP used in R21 (and RTS,S) vaccine constructs and the corresponding lack of the N-terminal JR. Heat maps illustrate the signal loss (OD) in four VAC065 volunteers when measured by indirect ELISA using NANP_6_ peptide or JR peptide (JR: KQPADGNPDPNANPNVDPN) after affinity chromatography and the depletion of NANP_6_-reactive IgG from day 84 plasma at pre-challenge (C_-1_) and post-challenge (C_+35_) time points. Bar graph illustrates Ig-Seq analysis of pre-challenge IgG in one volunteer (MA-1010).

To validate this at the molecular level, Ig-Seq analysis of pre-challenge IgG from one volunteer (MA-1010) using two separate pulldowns (NANP_6_ and JR) demonstrated that the JR-reactive repertoire overlapped with the NANP_6_-reactive repertoire lineage-for-lineage. Thus, in this volunteer, the JR-binding antibody lineages were a subset of the broader anti-NANP_6_ repertoire. Of the seven JR cross-reactive lineages identified in MA-1010 (meeting quality criteria and exceeding 0.5% relative abundance), there was diverse IGHV gene usage, with only one lineage employing an *IGHV3-30/33* family member (VH3-30). These seven lineages exhibited minimal somatic mutation in their IGHV gene segments (average 2.6%). The predominant lineage (VH3-7 at 47.5% relative abundance) had merely a single Asn52 mutation in its CDR-H2. Asn52 was previously identified as a key interacting residue in the dual-reactive monoclonal antibodies Ab668 and CIS42 targeting NANP/JR (Kisalu et al., 2018; Oyen et al., 2020).

Overall, these results suggest the NANP polypeptide of the central repeat region in the R21/Matrix-M vaccine is sufficient to instigate the generation and/or evolution of diverse, near-germline IgG antibodies that are cross-reactive with the N-terminal junction—even though the JR is absent from the vaccine construct.

### R21/Matrix-M vaccine-induced *IGHV3*-family antibodies show robust anti-sporozoite activity

From VAC053 immunized volunteers, a small panel of nine PfCSP plasma mAbs (VH:VL pairs) were isolated from day 63 (day 7 post-third vaccination) peripheral blood B cells. These nine plasma mAbs mapped with high confidence to proteomic CDR-H3 peptides of NANP_6_-reactive IgG lineages identified by Ig-Seq proteomic analysis (and circulating at >0.5% relative abundance). MAbs were expressed recombinantly as full-length IgG1 and first tested for NANP_6_ peptide-binding specificity by ELISA. Anti-sporozoite binding and inhibitory activity were also initially evaluated and compared with two previously characterized monoclonal antibodies derived from B cells of naturally exposed individuals in Africa (Triller et al., 2017); these two mAbs, 663 and 580, can inhibit Pf sporozoite traversal in hepatocytes *in vitro* and recognize transgenic *P. berghei* parasites expressing Pf CSP (PfCSP@PbCSP).

Three positive IgG plasma mAbs were prioritized for full, replicate downstream experimentation (**Table 2**). All three antibodies use *IGHV3-30/33* genes and represent a range of serological abundances, extremes of somatic mutation, and diverse expression of Ig light chains: MA1 (13% abundance; 1.6% SHM; *IGLV1-40*), MA6 (0.6%; 1.7%; *IGKV1-5*), and MA8 (19.6%; 8.2%; *IGKV1-16*). One negative *IGHV1-69* mAb (MA5) was also carried forward as a control reagent. All three *IGHV3-30/33* mAbs (MA1, MA6, and MA8) were NANP-specific and did not bind the JR peptide according to conventional ELISA and a novel flow cytometry assay (**Fig. S3** and data not shown).

**Table 2.**
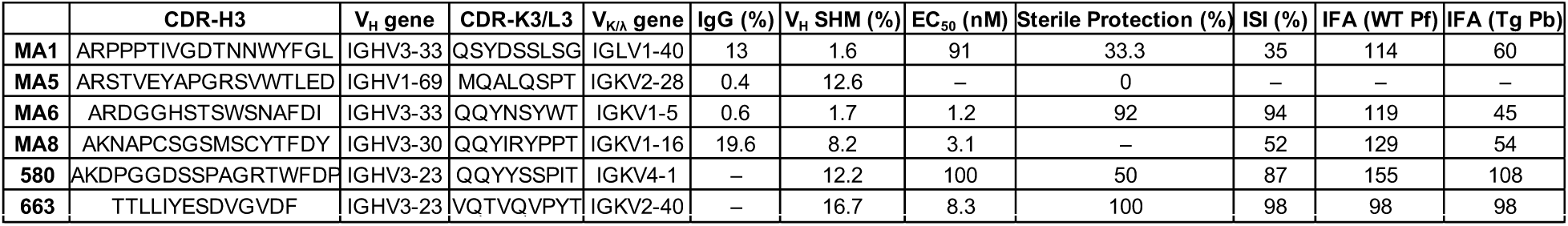
Summary of anti-NANP_6_ monoclonal antibodies used in this study. MA1, MA5, MA6, and MA8 were isolated from four different VAC065 volunteers. Control mAbs 580 and 663 previously reported by Triller et al., 2017. IgG (%): the percent relative abundance of the clone’s lineage in the total IgG anti-NANP_6_ repertoire SHM: somatic hypermutation at the nucleotide level ISI: inhibition of sporozoite invasion IFA: immunofluorescence assay

MAbs MA1, MA6, and MA8 bound strongly to wild-type 3D7 sporozoites in an immunofluorescence assay (IFA) and to transgenic (PfCSP@PbCSP) sporozoites (**Fig. 4 A**). The functional capacity of MA1, MA6, and MA8 was then assessed using an *in vitro* model of liver-stage development using cultured human hepatocytes to measure inhibition of sporozoite invasion (ISI) activity with PfCSP@PbCSP. All mAbs tested, including 580 and 663, were able to block hepatocyte invasion with MA1 and MA8 demonstrating a reduction in infection of 35% and 52%, respectively, while 580, 663, and MA6 all demonstrated more than 85% ISI activity (**Fig. 4 B**). To assess the ability of plasma antibodies induced by R21/Matrix-M to confer protection *in vivo,* we replicated the Triller et al. (2017) experimental scheme and included mAbs 580 and 663 as references. The model employs a transgenic strain of *P. berghei* (Pb), where the endogenous PbCSP is replaced with full-length PfCSP (PfCSP@PbCSP), enabling the assessment of PfCSP mAbs within the context of natural infection in a normal mouse, as rodents are permissive for Pb infection (Espinosa et al., 2017). Female C57BL/6 mice received 400 micrograms of antibody intraperitoneally and were challenged the following day with 5000 Pb-PfCSP sporozoites.

**Figure 4.**
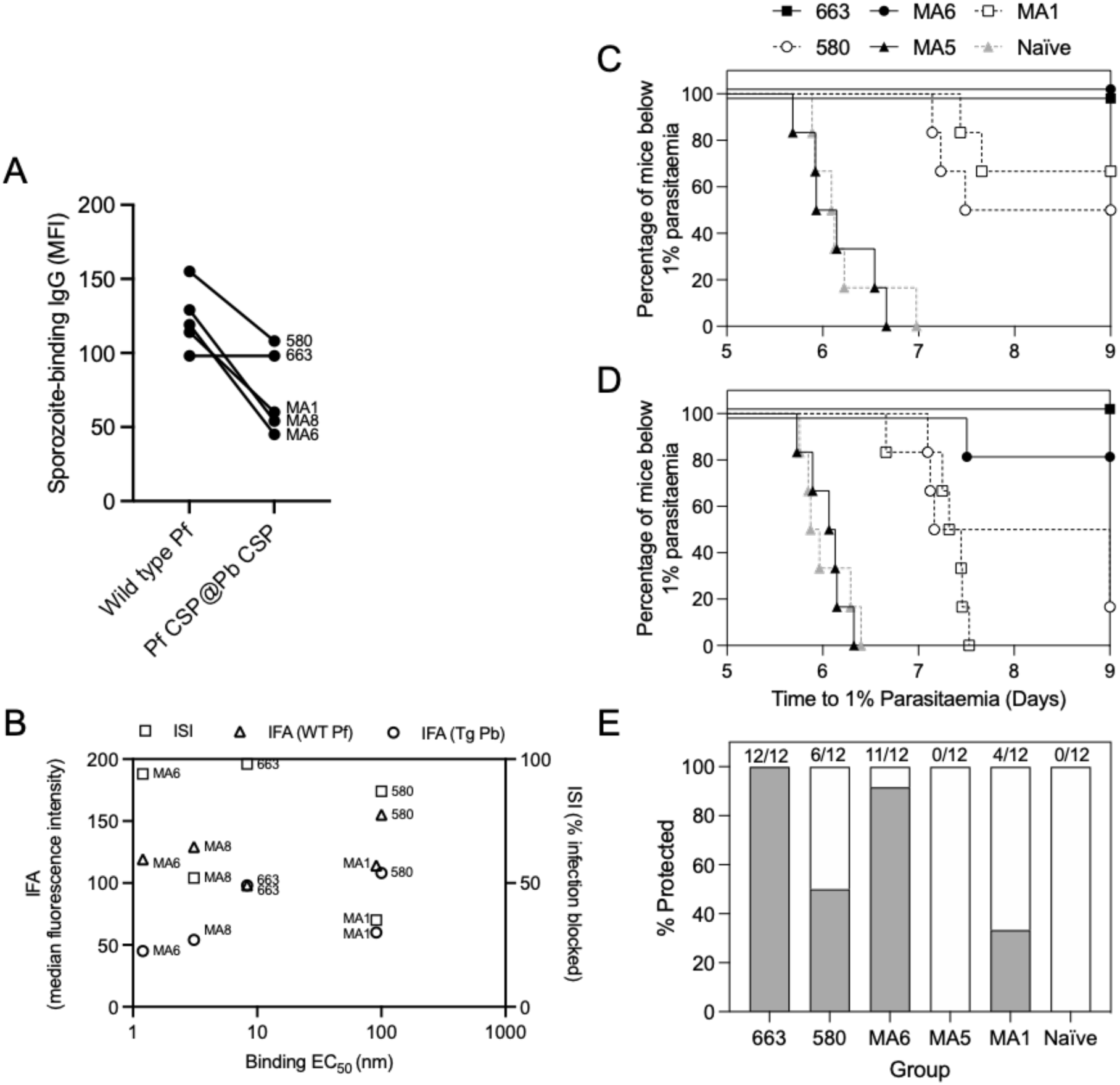
Monoclonal IgG plasma antibodies elicited by R21/Matrix-M vaccine provide sterile protection in mice. **(A)** Binding of mAbs to either wild-type 3D7 *Plasmodium falciparum* (WT Pf) or transgenic *P. berghei* expressing PfCSP (Tg Pb) measured by indirect immunofluorescence assay (IFA). (**B)** Relationship between either IFA or inhibition of sporozoite invasion (ISI) and NANP binding EC_50_. (**C– D).** C57BL/6J mice (n=6 per group) were injected intraperitoneally with 400 µg of monoclonal Ab on day −1. The following day mice were challenged with 5000 Pb-PfCSP *P. berghei* sporozoites administered by subcutaneous injection at the base of the tail. From 5 days post-sporozoite injection, mice were monitored by thin blood film for the development of blood-stage malaria. Parasitaemia was calculated daily and linear regression used to determine the time mice reached a threshold of 1% parasitaemia. Sterile protection was defined as any animal that remained parasite-free until day 9 after sporozoite administration. C and D are replicate experiments. (**E.** Summary of sterile protection for each mAb (grey bars represent the percent survival for each group over the 2 experiments).

Beginning five days after sporozoite injection, mice were monitored for protection against liver-stage malaria, indicated by reduction in time to develop blood-stage parasitaemia or complete absence of blood-stage parasites. In two independent experiments, MA1 and 580 sterilely protected 33% and 50% of mice, respectively, compared with naïve control mice, while MA5 as expected did not demonstrate protective immunity in this model (**Fig. 4 C–E**). MAb 580 demonstrated reduced efficacy (50% protection) in our model compared with 72% in the original experiment by Triller and colleagues. MA6 and 663 protected 92% and 100% of challenged mice, respectively, demonstrating potent inhibition of malaria infection. Of note, ISI appeared more closely related to *in vivo* protection than IFA or binding to NANP (**Fig. 4 B** and **E**).

### R21/Matrix-M vaccine can elicit IgG plasma antibodies structurally homologous to those induced in B cells after sporozoite exposure

The *IGHV3-33* germline-encoded tryptophan at position 52 in CDR-H2, which can also arise in some *IGHV3-30* alleles associated with anti-NANP responses, plays a crucial role in antigen binding, as previously observed in several PfCSP repeat-reactive antibodies identified following immunization with sporozoites or vaccination with RTS,S/AS01 (Pholcharee et al., 2020). To further explore the molecular nature of near-germline *IGHV3-33* antibodies generated by R21/Matrix-M vaccination, we selected monoclonal antibody MA6 for biophysical and structural studies (**Fig. 5**). Isothermal titration calorimetry experiments performed with the antigen-binding fragment (F_ab_) of MA6 and a representative peptide of the PfCSP repeat region (Ac-NPNANPNANPNA-NH_2_) resulted in a calculated binding affinity of ∼400 nM. Structural studies have clarified that anti-repeat binding motifs are in fact DPNA, NPNV, and NPNA, which result from the combination of major and minor repeats (Dyson et al., 1990; Pholcharee et al., 2020). The MA6 F_ab_ in complex with the peptide crystallized in space group P21 and diffracted X-rays to a resolution of 1.8 Å. The molecular replacement solution contained two F_ab_–peptide complexes in the asymmetric unit. Electron density for most of the peptide was well defined, allowing us to unambiguously place the first nine amino acids. The electron density for the remaining three C-terminal peptide residues was weaker and the corresponding residues could not be placed with confidence and were omitted from the model.

**Figure 5.**
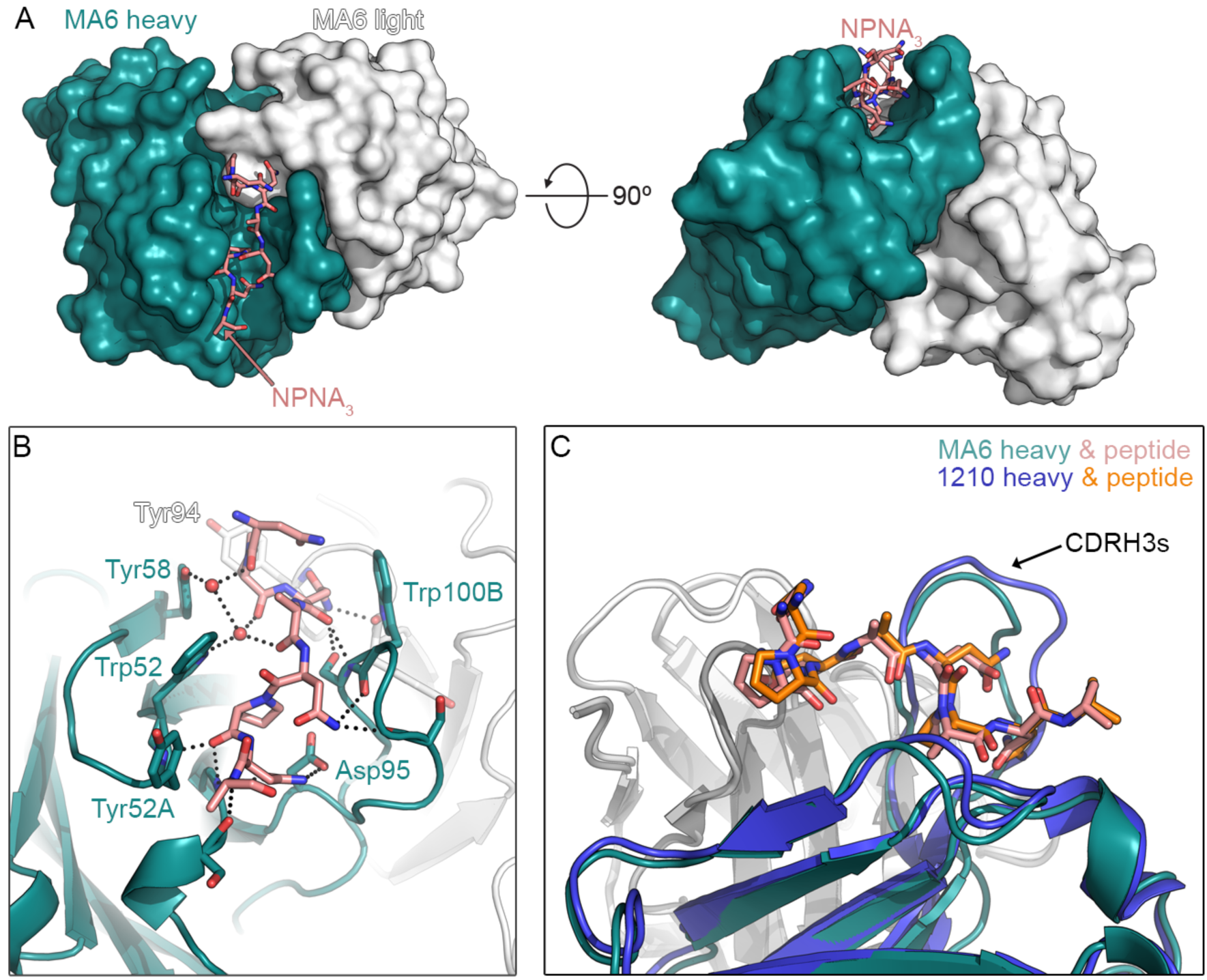
Elucidation of *IGHV3-33* antibody binding modes to the NANP polypeptide. Crystal co-structure of MA6 bound to NPNA_3_ peptide. (**A)** Overview of peptide in groove. The peptide lies within a groove formed primarily by the heavy chain, with CDR-H3 residues forming one side of the groove and CDR-H1 and -H2 residues forming the other side. (**B)** Specific contacts. Compared to other structures, the MA6 Trp52 sidechain is flipped such that the pyrrole ring of the indole, rather than the benzene ring, packs against Pro6 of the peptide. (**C)** Superposition with 1210 F_ab_. MA6 and 1210 bind the peptide in an almost identical conformation (RMSD of 0.23 Å for the first 8 C*α* atoms).

The structure revealed that the peptide lies within a groove formed primarily by the heavy chain, with CDR-H3 residues forming one side of the groove and CDR-H1 and -H2 residues forming the other side (**Fig. 5 A**). In total, there is 701 Å^2^ of buried surface area on the peptide, 406 Å^2^ buried on the heavy chain and 110 Å2 buried on the light chain, with the latter interface composed entirely of the CDR-L3. CDR-H3 residue Trp100B forms Van der Waals interactions with the peptide, as do CDR-H2 residues Trp52, Tyr52A, and Tyr58. On the CDR-L3, only the sidechains of Tyr94 and Trp96 substantially contribute to the binding interface with the peptide.

In addition to the hydrophobic interactions, numerous hydrogen bonds are formed between the peptide and antibody. Several are water-mediated, including those involving the sidechains of CDR-H2 residues Trp52 and Tyr58. The sidechains of CDR-H3 residues Asp95 and Ser100C both directly hydrogen bond to the peptide, but otherwise all remaining hydrogen bonds involve antibody main chain atoms. This allows for sequence variability in the antibody to be accommodated and likely explains the lack of CDR-H3 sequence conservation among similar antibodies.

Several structures of NANP_6_-reactive antibodies in complex with peptides have been determined, including antibodies 311 and 1210 (Imkeller et al., 2018; Murugan et al., 2018; Oyen et al., 2017). Of these, antibody 1210, which was obtained from a volunteer immunized with the Sanaria PfSPZ vaccine, is most like MA6, with 89% sequence identity between the heavy chains and 96% sequence identity between the light chains. Despite this overall high sequence similarity, the MA6 CDR-H3 is substantially different from the 1210 CDR-H3, with only 5 of 14 residues in common. Yet these antibodies bind the peptide in an almost identical conformation, resulting in an RMSD of 0.23 Å for the first 8 C_α_ atoms. X-ray crystallography of CSP-binding antibodies has previously revealed a key structural requirement for Trp52 in the CDR-H2 of the *IGHV3-33* gene element; this feature is conserved in human mAbs 1210, MGG4, and 311 whether elicited by live sporozoites or RTS,S immunizations, as has been reviewed (Oyen et al., 2017; Pholcharee et al., 2021; Scally and Julien, 2018). Compared to other structures, the MA6 Trp52 sidechain is flipped such that the pyrrole ring of the indole, rather than the benzene ring, packs against Pro6 of the peptide. Collectively, the structure of MA6 in complex with a NPNA_3_ peptide and its comparison to the peptide-bound 1210 structure provide a molecular basis for the dominance of near-germline *IGHV3-33* NANP_6_-reactive antibodies and their ability to accommodate diversity in their CDR-H3s.

## DISCUSSION

The field of malaria vaccines has made significant progress, highlighted by the WHO’s recommendation of two vaccines for children in regions with moderate to high transmission of Plasmodium falciparum. The WHO expects that the introduction of the PfCSP-based VLP vaccine, R21/Matrix-M, alongside the ongoing rollout of the first malaria vaccine, RTS,S, will help meet global demand. The robust efficacy rates observed with R21/Matrix-M at 12 months in areas with year-round malaria transmission (Datoo et al., 2024), along with promising preliminary data at 42 months in seasonal transmission sites (Natama et al., 2023), indicate that R21 is an efficacious vaccine with significant potential to reduce the burden of malaria, comparable to the expected impact of RTS,S. Our molecular-level analysis of the R21-generated IgG plasma antibody repertoire—showing global similarity to previously reported RTS,S B-cell repertoires and validating the robust potency of these antibodies—support this conclusion.

A significant finding in this study is the dominance of NANP repeat-specific antibodies with minimal somatic hypermutation in the plasma IgG repertoire induced by the R21/Matrix-M malaria vaccine. Like RTS,S-induced B-cell repertoires (Oyen et al., 2017; Williams et al., 2024), the IgG plasma repertoires elicited by R21/Matrix-M also are predominantly built upon germline or near-germline *IGHV3*-family genes (*IGHV3-30/3-33*). Through the cloning, functional, and structural analysis of prevalent IgG anti-NANP antibodies we demonstrated how the R21-generated plasma antibody, MA6, inhibits sporozoite cell invasion in cell culture, curtails parasitemia, protects in a transgenic mouse model, and mediates NANP-recognition through a canonical CDR-H2 Trp52 structural motif encoded by *IGHV3-33*. Since the MA6 VH region differs from the germline *IGHV3-33* gene segment by only two amino acids in its framework 2 region— neither of which interacts with NPNA_3_ in the F_ab_-peptide crystal structure—we speculate that MA6 originated from the naïve B-cell pool and may have seeded the IgG-secreting plasma cell repertoire via a non-canonical maturation pathway. Our data suggest that repeat-specific antibodies targeting the NANP region may not undergo the extensive somatic hypermutation typically seen in more mature antibody responses, instead favoring an innate-like, early-stage antibody pool that is rapidly mobilized following vaccination. This speculation aligns with both recent and past observations based on RTS,S and whole sporozoite immunizations (Murugan et al., 2018; Murugan et al., 2020; Williams et al., 2024). Furthermore, the MA6 VL region is encoded by unmutated *IGKV1-5* and contains an eight amino-acid long CDR-K3, akin to the “public” NANP-reactive “VH3-33/Vk1-5/KCDR3:8” antibody class repeatedly observed in CSP B-cell repertoires (Murugan et al., 2018; Tan et al., 2018; Williams et al., 2024). Our data thus establish a connection between CSP/NANP immune responses in the naïve B-cell compartment and their evolution into secreted, prevalent IgG circulating in the blood.

Notably, our study also demonstrated that controlled human malaria infection (CHMI) challenge does not significantly boost the plasma IgG response. Despite the challenge, the quantity and composition of the NANP-reactive antibody repertoire remained stable. This finding, although counterintuitive, highlights the robust and static nature of the antibody response generated by the R21/Matrix-M vaccine. Since the elicitation of neutralizing IgG antibodies correlates with protection by vaccination for most licensed vaccines (Plotkin, 2010), and IgG anti-CSP titers meet the criteria for an immunological correlate of protection for the R21/Matrix-M vaccine (Schmit et al., 2024), we propose that the molecular profile of circulating antibodies elicited by R21/Matrix-M, as identified by Ig-Seq proteomic analysis, must reflect this fundamental biology. In this context, the temporal stability and static composition of R21-generated antibodies up to five weeks after a sporozoite challenge infection underscore the vaccine’s ability to elicit an effective sporozoite-neutralizing response. Among the IgG antibodies involved in this response, we found that R21/Matrix-M generates antibodies targeting the CSP junctional region, in addition to the CSP NANP repeat region encoded by the vaccine immunogen, representing a subset of the broader anti-NANP_6_ repertoire. Although further studies are needed, these cross-reactive antibodies may belong to the same class as the prophylactic mAbs CIS43 and L9, which have demonstrated high efficacy in advanced clinical field trials (Kayentao et al., 2022; Kayentao et al., 2024).

Both currently recommended malaria vaccines are rapidly being introduced throughout sub-Saharan Africa. In addition, efforts are underway to enhance existing vaccine efficacy by developing attenuated whole sporozoite, blood stage, and transmission-blocking vaccine candidates. There is consensus that a multistage vaccine approach, and perhaps their integration with monoclonal antibodies, will be required for the elimination of malaria. Research indicates that a dual approach (combining monoclonal antibodies and vaccines) can provide synergistic effects, offering higher levels of protection than either strategy alone (Wang et al., 2021). Likewise, integrating the pre-erythrocytic R21/Matrix-M vaccine with blood-stage candidates, like the promising strain-transcendent RH5/Matrix-M (Silk et al., 2024), could provide even stronger protection. However, this will necessitate the development of clinical plans to test combination vaccines targeting both pre-erythrocytic and blood stages (PATH, 2024).

The application of Ig-Seq proteomics to analyze circulating antibodies produced by the R21/Matrix-M malaria vaccine at the molecular level demonstrates the power of this technique for evaluating pre-erythrocytic vaccines. Molecular deconvolution of the serological immune response through LC–MS/MS-based approaches could also shed light on long-standing enigmas, such as dysregulated B-cell activation (“Plasmodium jamming” (Klein, 1982)), hypergammaglobulinemia, and autoantibody production during acute infection (Crompton et al., 2014; Donati et al., 2004); the heightened presence of circulating immune complexes associated with cerebral malaria and severe malarial anemia (Pleass, 2009); and the dynamic interactions between neutralizing and non-neutralizing antibodies during blood stage malaria (Barrett et al., 2024).

## MATERIALS & METHODS

### Plasma and PBMC samples

Plasma and PBMC samples were collected from ten healthy malaria-naïve United Kingdom adult volunteers. For four volunteers, blood was drawn seven days (day 63) following their third and final (day 56) vaccination with R21/Matrix-M (VAC053; clinicaltrials.gov identifier NCT02572388). For six volunteers, blood was drawn 28 days (day 84) following their third and final (day 56) vaccination. (VAC065; clinicaltrials.gov identifier NCT02905019). The six VAC065 volunteers also had blood drawn 35 days (day 112) following a day 85 challenge infection by mosquito-bite transfer of *Plasmodium falciparum* 3D7 sporozoites. Both clinical trials were conducted by the Jenner Institute at the University of Oxford.

### High-throughput sequencing of BCR transcripts

Frozen PBMCs were thawed at 37°C, resuspended in RPMI-1640 (Fisher) supplemented with 10% Fetal Bovine Plasma (FBS) and 20 U/mL deoxyribonuclease I (Roche), concentrated by centrifugation (300 x g for 15 min at 20°C), and allowed to recover in 4 mL media at 37°C for 30 min. The cells were then split: half were dedicated to BCR heavy chain transcript sequencing, and half were used to identify the native VHVL repertoire.

### VH Sequencing

Cells were diluted with 10 mL of cold buffer (phosphate buffered saline supplemented with 0.5% BSA and 2 mM EDTA) and pelleted via centrifugation (300 x g for 15 min at 4°C). The supernatant was decanted, and the cells were resuspended in 1 mL of TRIzol^TM^ reagent (Thermo Fisher). RNA was extracted using RNeasy (Qiagen), and first strand cDNA was synthesized from 500 ng total RNA using SuperScript IV (Invitrogen). IgG, IgA, and IgM heavy chain repertoires were amplified using a multiplex primer set (Ippolito et al., 2012) and sequenced by 2×300 paired-end Illumina MiSeq.

### VH:VL paired sequencing

Total B cells were isolated from PBMCs using the Human Memory B Cell Isolation Kit (Miltenyi Biotec) with an LD column according to the manufacturer’s instructions. B cells were then coemulsified with poly-dT magnetic beads and lysis buffer using a custom-designed flow-focusing device as described (McDaniel et al., 2016). VH and VL amplicons from single B cells were amplified, stitched together using overlap-extension RT-PCR, and sequenced by 2×300 Illumina MiSeq.

### Isolation of NANP_6_-reactive antibodies from plasma

1-2 mL plasma from each volunteer was purified with Protein G Plus agarose affinity chromatography (ThermoFisher Scientific), and IgG F(ab′)_2_ fragments were generated with IdeS at a 1:50 ratio at 37°C for 2 h. NANP_6_-reactive F(ab′)_2_ fragments were isolated by affinity chromatography using NeutrAvidin resin (Pierce) coupled to biotinylated NANP_6_ peptide (ABclonal), and eluted in 100 mM Glycine (pH 2.7).

### Sample preparation and LC-MS/MS bioinformatic analysis

For each sample, the flow through (nonreactive to NANP_6_) and NANP_6_-reactive elution were separately denatured in 1:1 (v/v) trifluoroethanol, reduced with 5 µL dithiothreitol (110 mM) at 55°C for 45 min, and alkylated with 32 mM iodoacetamide by incubating in the dark at RT for 30 min. Samples were diluted 10x into 50 mM Tris (pH 8.0) and digested with trypsin (1:10 trypsin/protein) for 3 hours at 37°C. Formic acid (0.1%) was used to quench the reaction. The solution was then concentrated, desalted using a C18 Hypersep SpinTip (Thermo) according to the manufacturer’s instructions, and submitted for LC-MS/MS.

Samples were submitted to the UT Austin CBRS Proteomics Facility for protein identification by LC-MS/MS using the Dionex Ultimate 3000 RSLCnano LC coupled to the Thermo Orbitrap Fusion. A 2 cm long x 75 µm I.D. C18 trap column was followed by a 75 µm I.D. x 25 cm long analytical column packed with C18 3 µm material (Thermo Acclaim PepMap 100). The FT-MS resolution was set to 120,000, and 3-sec cycle time MS/MS were acquired in ion trap mode.

SEQUEST (Proteome Discoverer 1.4, Thermo Scientific) with previously described settings (Williams et al., 2017) was used to search the spectra against a patient-specific protein database constructed from the full-length VH and VL sequences, Ensembl human protein-coding sequences, and common contaminants (maxquant.org). PSMs were filtered with Percolator (Proteome Discoverer 1.4) at a false discovery rate of <1% and filtered for average mass deviations <1.5 parts per million. Peptides mapping to the CDR-H3 region of unique antibody lineages were grouped, and the relative abundances of the corresponding peptide matches were determined by the sum of the extracted ion chromatogram (XIC) peak areas of the respective precursor ions, as previously described (Lavinder et al., 2014; Lee et al., 2016; Voss et al., 2021).

### Antibody expression and purification

We chose monoclonal antibodies (mAbs) that demonstrated high-confidence identification of CDR-H3 peptides in plasma via LC-MS/MS, evidenced by high abundance (indicated by the XIC area) and extensive peptide coverage of the VH, particularly in the hypervariable CDR regions. Selected antibody sequences were purchased from Twist Bioscience as gene fragments cloned into a customized pcDNA3.4 vector (Invitrogen) containing human IgG1 Fc regions, IgK1 Fc regions, or IgL2 Fc regions. VH and VL plasmids were transfected into 30 mL cultures of Expi293F cells (Invitrogen) at a 1:2 ratio and incubated at 37°C and 8% CO_2_ for 7 days. The supernatant containing secreted antibodies was collected following centrifugation (1000 g for 10 min at 4°C), neutralized, and filtered. Antibodies were isolated using Protein G Plus agarose (ThermoFisher Scientific) affinity chromatography, washed with 20 column volumes of PBS, eluted with 100 mM glycine-HCl pH 2.7, and neutralized with 1 M Tris-HCl pH 8.0. The antibodies were then concentrated, and buffer exchanged into PBS using 10,000 MWCO Vivaspin centrifugal spin columns (Sartorius).

### Protein expression and purification for crystallization

The amino acid sequences for MA6 heavy and light chain variable domains were codon optimized and synthesized as gBlock gene fragments (Integrated DNA Technologies). Gene fragments were cloned into the mammalian expression plasmid pVRC8400 with an HRV 3C protease cleavage site inserted within the hinge region of the heavy chain sequence. Plasmids were used to co-transfect 0.5 L FreeStyle™ 293-F cells (Invitrogen) with 0.167 mg heavy chain and 0.083 mg light chain DNA using polyethylenimine 25K (Polysciences, Inc.). Six days after transfection, cells were harvested and centrifuged at 5,400xg before supernatant containing soluble protein was concentrated and buffer exchanged into PBS using a 30 kDa tangential flow filtration cassette (PALL). Sample was then passed over Pierce™ Protein A Plus agarose resin (Thermo Scientific) in a gravity flow column. Eluate was processed with HRV 3C protease and passed over a gravity flow column packed with CaptureSelect™ IgG-CH1 resin to purify the MA6 antigen binding fragment (F_ab_). A purification polishing step was then performed by passing the eluate over a Superdex 200 10/300 GL gel filtration column (Cytiva) in crystallization running buffer (2 mM Tris pH 8.0, 200 mM NaCl). F_ab_ peak fractions were concentrated to 11.88 mg/mL using an Amicon® 30,000 MWCO centrifugal spin column (EMD Millipore). NPNA_3_ peptide (ABclonal) was solubilized in crystallization buffer (2 mM Tris pH 8.0, 200 mM NaCl) to a stock concentration of 500 µM.

### Crystallization and data collection of MA6 in complex with NPNA_3_

F_ab_ and NPNA_3_ peptide were combined at a 1:12 molar ratio and concentrated to 12.5 mg/mL. Crystals were produced using the sitting-drop vapor diffusion method by mixing 0.1 µL of MA6 F_ab_-NPNA_3_ complex with 0.05 µL of reservoir solution containing 27% PEG 3350 and 3% MPD. The crystal was flash-cooled in liquid nitrogen directly from the crystallization drop. Diffraction data were collected to 1.8 Å from a single crystal at SBC beamline 19ID (Advanced Photon Source, Argonne National Laboratory).

### Structure determination and refinement

Data were indexed and integrated in iMOSFLM (Battye et al., 2011), before being merged and scaled using Aimless (Evans and Murshudov, 2013). Molecular replacement was performed in Phaser (McCoy et al., 2007), and the model was subjected to multiple rounds of building and refinement in Coot (Emsley and Cowtan, 2004) and PHENIX (Adams et al., 2002), respectively. Data collection and refinement statistics can be found in Table 3. SBGrid (Morin et al., 2013) compiles all the crystallography processing and refinement programs used in this study.

**Table 3.** MA6 F_ab_ + NPNA_3_ data collection and refinement statistics.

| PDB ID | 9CR9 |
| --- | --- |
| **Data collection** |  |
| Wavelength (Å) | 0.9795 |
| Resolution range | 47.72 1.8 (1.864 1.8) |
| Space group | *P* 1 2<sub>1</sub> 1 |
| Unit cell dimensions |  |
| *a*, *b*, *c* (Å) | 46.6, 61.0, 153.0 |
| $\alpha$, $\beta$, $\gamma$ (°) | 90, 90, 90 |
| Total reflections | 137,629 (13,558) |
| Unique reflections | 74,608 (7,426) |
| Multiplicity | 1.8 (1.8) |
| Completeness (%) | 93.3 (93.1) |
| *I*/σ*I* | 5.93 (2.14) |
| Wilson B-factor | 17.1 |
| *R*<sub>merge</sub> | 0.077 (0.4335) |
| CC<sub>1/2</sub> | 0.941 (0.626) |
| **Refinement** |  |
| Unique reflections | 74,522 (7,401) |
| *R*<sub>work</sub> | 0.1853 (0.2749) |
| *R*<sub>free</sub> | 0.2153 (0.3039) |
| Number of non-hydrogen atoms | 7,814 |
| macromolecules | 6,652 |
| ligands | 0 |
| solvent | 1,162 |
| Protein residues | 872 |
| RMSD bond lengths (Å) | 0.008 |
| RMSD bond angles (°) | 0.96 |
| Ramachandran favored (%) | 98.36 |
| Ramachandran allowed (%) | 1.64 |
| Ramachandran outliers (%) | 0.00 |
| Rotamer outliers (%) | 0.00 |
| Clashscore | 4.05 |
| Average B-factor (Å<sup>2</sup>) | 23.9 |
| macromolecules | 22.1 |
| solvent | 34.1 |

Statistics for the highest-resolution shell are shown in parentheses.

### Immunofluorescence assay (IFA)

Chambered microscope slides coated with either wild-type 3D7 *Plasmodium falciparum* or transgenic *P. berghei* expressing PfCSP sporozoites were stored at −80°C until use. Slides were brought to room temperature (RT) and then fixed for 15 minutes in 4% paraformaldehyde. After washing twice in PBS for 5 minutes, slides were blocked for 1 hour in casein. Slides were washed as before and 10 µL of serum sample diluted 1:100 in casein was added to each well. Slides were incubated for 30 minutes at RT in a humidity chamber then wells were individually washed with PBS three times for 5 minutes. Secondary antibody (anti-IgG-Alexafluor488) was diluted 1:800 in casein and 15 µL was added to each well. Slides were incubated for 30-45 minutes in a humidity chamber at RT protected from light. After a final wash, slides were rinsed in distilled water and left to dry before mounting with DAPI-containing media. Slides were set overnight at 4°C before being examined under a Lecia DMI3000 B microscope. Images were captured in QCapturePro software (Surrey, BC, Canada) using brightfield illumination, GFP and DAPI filters at set exposure levels. ImageJ software was used to measure the median fluorescent intensity for 5 sporozoites in each well and an average was taken.

### Inhibition of sporozoite invasion (ISI) assay

Inhibition of sporozoite invasion was assessed as previously described (Bowyer et al., 2018; Rodríguez-Galán et al., 2017). Human hepatoma cells (HC04 or Huh7) cultured in RPMI with 10% FBS were added to 96-well plates at 30,000 cells/well and left to settle overnight at 37°C, 5% CO2. Viable GFP-labelled *Plasmodium berghei* sporozoites expressing *P. falciparum* CSP at the P. berghei CSP locus (P. berghei PfCSP@PbCSP) were obtained by dissecting infected *Anopholes stephensi* mosquitoes. Dissected salivary glands were pooled into RPMI 1640. Culture medium was aspirated from the hepatoma cells, then 100 µL of serum diluted 1:5 in RPMI with 10% FBS and 100 µL of sporozoite solution (10,000 sporozoites) were added to each well, giving a 10% final serum concentration. Samples were tested in duplicate: “hepatoma only” wells and positive control wells containing hepatoma cells and sporozoites, but no serum, were included. Baseline (pre-vaccination) and post-vaccination samples were run for each volunteer tested. After incubation for 20-26h at 37°C, media was aspirated and plates were washed with 90 µL/well DPBS. Cells were trypsinized, re-suspended in 65 µL DPBS with 1% bovine serum albumin (BSA) and acquired immediately using a BD LSR II and FACSDiva v6.2. DAPI stain was added to each sample just before acquisition. Data was analysed in FlowJo software v10.6 (TreeStar Inc., Ashland, Oregon).

### Protection study in mice

Mice were used in accordance with the UK Animals (Scientific Procedures) Act under project license number P9804B4F1 granted by the UK Home Office following ethical review by the University of Oxford Animal Welfare and Ethical Review Board (AWERB). Animals were group housed in IVCs under SPF conditions, with constant temperature and humidity with lighting on a 13:11 (7am to 8pm) light-dark cycle. Each mouse received 400 µg of mAb intraperitoneal injection (i.p.) the day before challenge with 5000 Pb-PfCSP sporozoites.

Plasmodium berghei sporozoites (spz) expressing Plasmodium falciparum CSP under the control of the PbCSP promoter (Pb-PfCSP) (Rodríguez-Galán et al., 2017) were isolated from salivary glands of female *Anopheles stephensi* mosquitoes around 21 days after feeding on a Pb-PfCSP blood stage infected donor mouse. Salivary glands were homogenised, sporozoites counted under phase contrast microscopy and 5000 Pb-PfCSP sporozoites injected sub-cutaneously (s.c.) into recipient mice. Mice were monitored daily from day 5 onwards by taking a thin blood film and staining with 5% Giemsa (Sigma Aldrich) to screen for the presence of schizonts within the red blood cells. Parasitaemia was calculated as the percentage of infected red blood cells per microscope field (100x objective), with at least 5 fields counted per mouse per day. Using linear regression, the time to 1% parasitaemia was calculated based on the y-intercept and slope of the line.

### Statistical analysis

Non-parametric Mann–Whitney U-test was used to compare two groups. Two-tailed Kruskal-Wallis test followed by Dunn’s multiple comparisons test was used to compare groups of three or more. Kaplan–Meier survival analysis was tested by log-rank test. Sample sizes for each experiment were based on a power calculation and/or previous experience of the mouse models. Data analysis and visualization were performed using GraphPad Prism, Microsoft Excel, Microsoft PowerPoint, Adobe Illustrator, Immunarch, and R Studio. P-values < 0.05 were considered statistically significant.

## Author Contributions

Writing - Original Draft: J.R. McDaniel, W.N. Voss, S.A. Rush, K.J. Ewer, and G.C. Ippolito; Writing - Review & Editing: S. Gregory, C.R. King, J.R. McDaniel, J.S. McLellan, A.V.S. Hill, G. Georgiou, W.N. Voss, K.J. Ewer, and G.C. Ippolito; Investigation: J.R. McDaniel and W.N. Voss performed the BCR-Seq and Ig-Seq experiments. S.A. Rush performed the X-ray crystallography experiments. G. Bowyer, A.J. Spencer, D. Bellamy, and M. Ulaszewska performed the inhibition of sporozoite invasion experiments, mouse protection experiments, and immunofluorescence assays. J. Goike performed the flow cytometry junctional peptide binding experiment; Formal analysis: J.R. McDaniel, W.N. Voss, G. Bowyer, S.A. Rush, A.J. Spencer, D. Bellamy, J. Goike, J. McLellan, K.J. Ewer, G.C. Ippolito; Visualization: J.R. McDaniel, W.N. Voss, S.A. Rush, K.J. Ewer, and G.C. Ippolito.

## ACKNOWLEDGEMENTS

This work was funded by PATH’s Center for Vaccine Innovation and Access, through a grant awarded to PATH by the Bill & Melinda Gates Foundation (INV-007217), under a collaborative agreement with The Jenner Institute at the University of Oxford (K.J.E.) and The University of Texas at Austin (G.C.I.). This work was partly funded by Welch Foundation grant number F-0003-19620604 (J.S.M.). Next-generation sequencing was performed by the Genomic Sequencing and Analysis Facility at UT Austin, Center for Biomedical Research Support, RRID#: SCR_021713. Mass spectrometry was performed by the Biological Mass Spectrometry Facility at UT Austin. The authors acknowledge the Texas Advanced Computing Center (TACC) at The University of Texas at Austin for providing HPC resources that have contributed to the research results reported within this paper.

---

**Figure S1.**
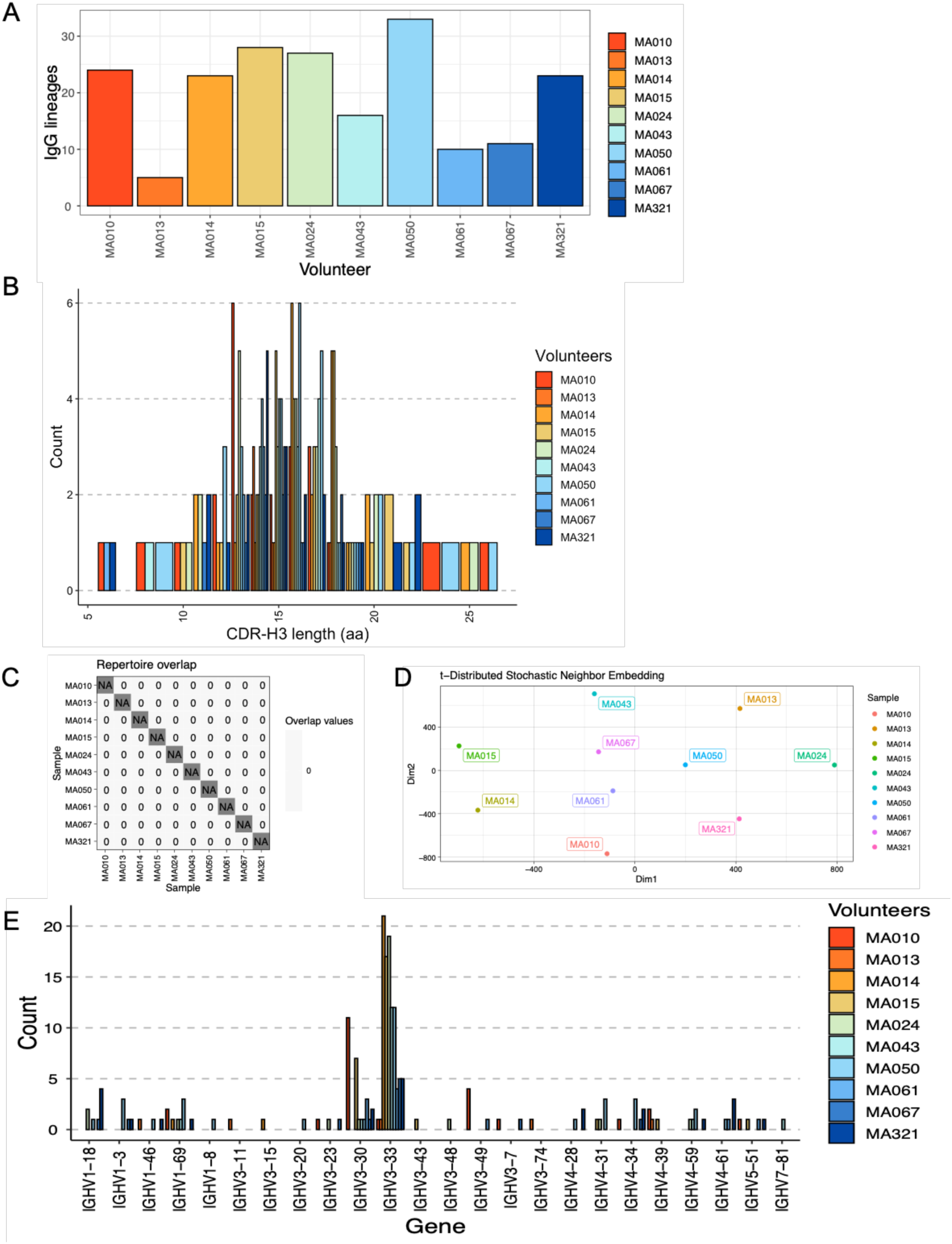
Profile of NANP_6_-reactive IgG repertoires across all ten volunteers post-vaccination. **(A)** Number of IgG lineages per volunteer. **(B)** Frequency of CDR-H3 lengths for all lineages. **(C)** Checkerboard graphic illustrating no shared CDR-H3s (no overlap) across volunteers. **(D)** Dissimilarity of VH repertoires according to t-distributed stochastic neighbor embedding (tSNE) analysis. **(E)** Frequency of IGHV gene use by volunteer.

**Figure S2.**
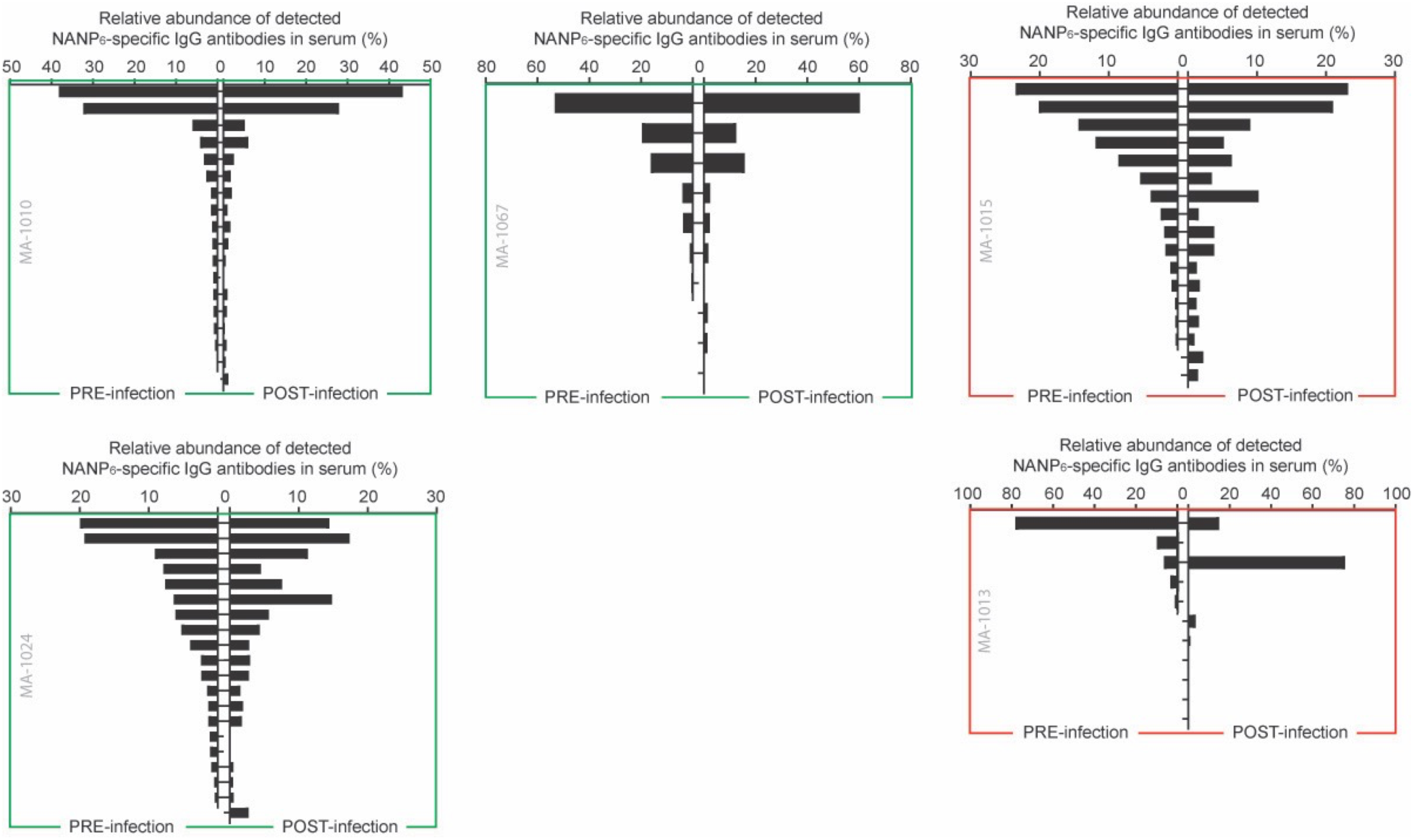
Static composition and high correlation between NANP_6_ pre-challenge and post-challenge IgG repertoires among VAC065 volunteers. Green outline indicates the volunteer was protected from infection; red that they were not protected.

**Figure S3.**
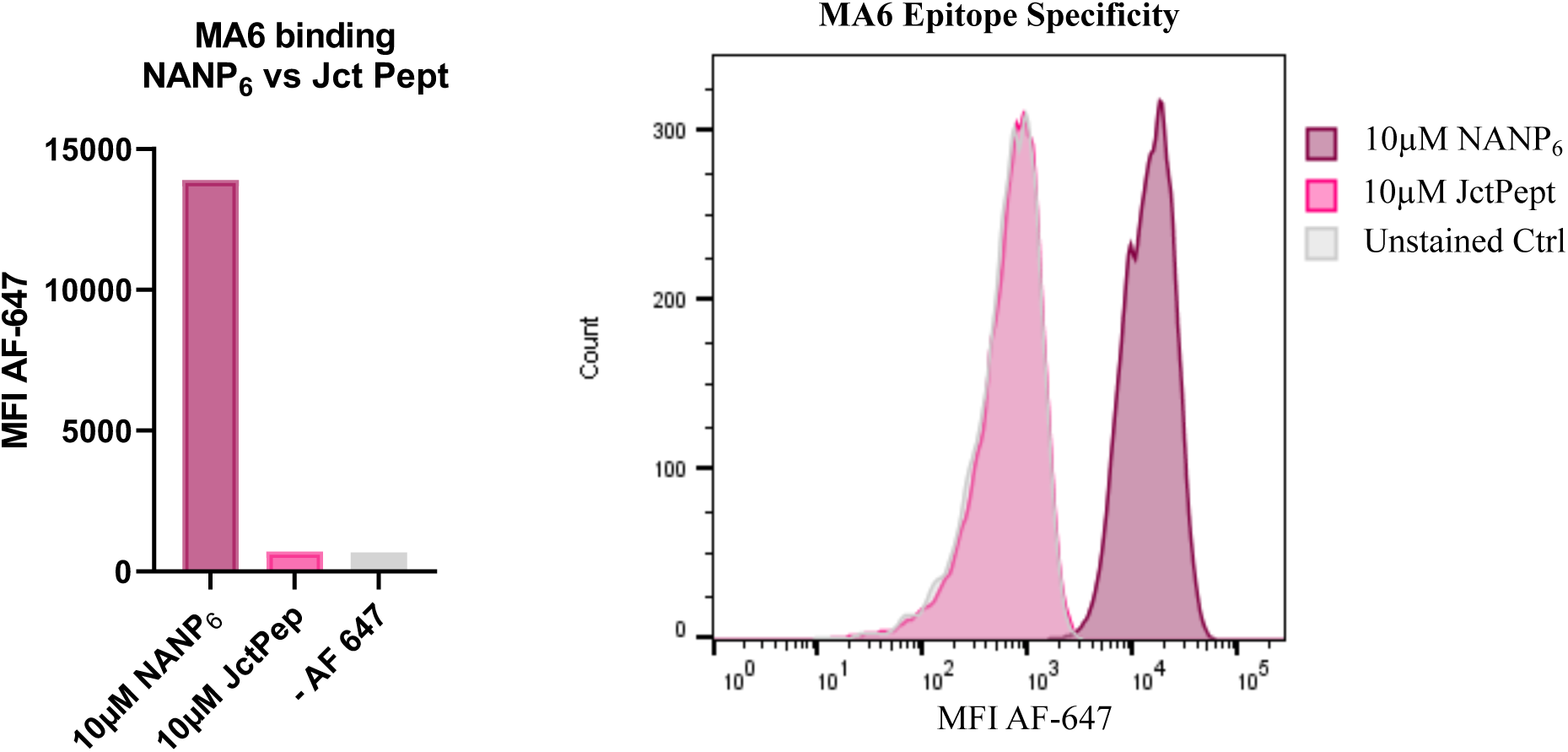
MA6 Epitope Specificity. MA6 Fab-fragment displaying yeast were incubated with 10 µM biotinylated NANP_6_ or junctional region peptide (Jct Pept: KQPADGNPDPNANPNVDPN) and subsequently labeled with streptavidin-conjugated Alexa-Fluor 647 to detect binding via flow cytometry.

